# An Analysis of the Value-Added of Antibiogram Subgroup Stratification

**DOI:** 10.1101/2024.08.29.610362

**Authors:** Connie T. Y. Xie, Samantha Martinez, Ceylon V. Simon, Susan M. Poutanen

## Abstract

**Background:** Stratified antibiograms are recommended to guide empiric clinical treatment. However, which strata to focus on, the limited number of isolates in identified strata, and the heavy associated workload all pose challenges. This study compares differences in antibiotic susceptibility between a hospital-wide, all-specimens antibiogram and stratified antibiograms in order to identify the value-added of antibiogram stratification.

**Method:** Antibiotic susceptibility of bacterial isolates from 2021 at a quaternary-care academic hospital was obtained from published hospital-wide and unit- and specimen-specific stratified antibiograms. Differences in percent susceptibility by organism and drug between the hospital-wide and stratified antibiograms were calculated. Weighted averages of the difference in percent susceptibility were calculated for each stratified antibiogram compared to the hospital-wide antibiogram and unit-wide antibiograms. Differences were shown through heat maps.

**Results:** When compared to a hospital-wide, all-specimens antibiogram, the emergency department (ED) antibiogram showed higher susceptibility, whereas intensive care unit (ICU) and, particularly, transplant unit (TR) antibiograms exhibited reduced susceptibility. Compared to ward level antibiograms, further stratification within each ward to specimen-specific (syndromic) antibiograms revealed additional differences. In ED, urine and respiratory-stratified antibiograms had lower susceptibility and blood had higher susceptibility. Compared to ward-specific antibiograms, in ICU, all specimen-stratified antibiograms had lower susceptibility and in transplant, antibiograms for all specimens but urine had lower susceptibility.

**Conclusion:** Using a hospital-wide all-specimens antibiogram may both overcall and under call susceptibility leading to poor empiric antimicrobial choices. Specimen-specific antibiograms stratified by unit best inform empiric therapy for specific populations.

**Summary:** Using a hospital-wide all-specimens antibiogram may both overcall and under call susceptibility leading to poor empiric antimicrobial choices. There is value-added to providing combined unit-specific and specimen-specific antibiograms to help best inform empiric therapy for specific populations.

## Introduction

Rising antimicrobial resistance poses an ongoing challenge for both empiric and definitive treatment of infections [1]. The Infectious Diseases Society of America (IDSA) recommends compilation of antibiograms as a way to allow for institutional surveillance of antimicrobial susceptibility and to serve as an empirical guide to antimicrobial treatment [2]. From clinical susceptibility tests, average susceptibility percentage of specific organisms-antimicrobial pairings are calculated to track changes in antimicrobial susceptibility. The Clinical and Laboratory Standards Institute (CLSI) publishes consensus guidelines (M39) on the methods of developing antibiograms to ensure their accuracy and comparability. The M39 outline methods regarding the collection, analysis and presentation of data [3]. Basic recommendations include that: antibiogram reports should be presented at least annually; only diagnostic isolates should be included; only the first isolate of a species per patient per analysis period should be included; and reported species should have ≥30 isolates [3]. The CLSI M39 document and IDSA guidelines suggests the use of stratified antibiograms over non-stratified to improve empiric antibiotic therapy in specific populations ([2] suggesting that a hospital-wide antibiogram may conceal differences in susceptibility across different healthcare parameters within a single institution, such as the hospital unit, the infection site, and patient population. The concern is that this can ultimately affect optimal patient treatment as well as tracking emerging antimicrobial resistance patterns that are relevant to certain settings.

Indeed, the aggregation of susceptibility data in hospital-wide antibiograms has been shown to be potentially misleading due to the masking of trends in specific patient settings. To date, comparison of hospital-wide to stratified antibiograms are largely focused on categories specific to hospital unit [4–12] with an emphasis on ICU differences, while some have also looked at impact of stratification by infection location [13], and patient characteristics [13–17]. Most studies have focused on investigating an a priori identified specific strata as opposed to looking at hospital wide differences across many strata.

Our laboratory serves a number of academic acute care and non-acute care health care facilities in the Greater Toronto Area. Our laboratory publishes hospital-wide antibiograms annually along with specific-specific antibiograms stratified by unit for all acute care hospital clients [18]. The challenges of creating these stratified antibiograms include knowing which strata to focus on, the limited number of isolates in stratified antibiograms, and the heavy workload associated with data clean-out and manipulation associated with the additional analyses required. Given the amount of data and its presentation as distinct antibiograms, it is difficult to determine the value-added of each stratified antibiogram. Using one of our larger acute care hospital’s data, the purpose of this study is to compare differences in antibiotic susceptibility between hospital-wide, all-specimens antibiogram and stratified antibiograms in order to identify the value added of subgroup stratification across all strata.

## Methods

Antibiotic susceptibility of bacterial isolates from 2021 of a quaternary-care academic hospital served by our laboratory was obtained from published hospital-wide and unit- and specimen-specific stratified antibiograms [18]. Susceptibility testing and antibiogram generation was performed in accordance with CLSI with the exception of including data for species with less than 30 organisms per strata but greater than 10 in order to be able to include all subgroup analyses. Differences in percent susceptibility by organism and antimicrobial between the reference antibiogram and stratified antibiograms were calculated (Appendices 1-5). Weighted averages (WA) of the change in percent susceptibility, excluding coagulase-negative staphylococci, was calculated for each stratified antibiogram using the number of organisms on the stratified antibiogram as the weight. Heat maps representing these values were created. Excel and R Statistical Software (4.2.1., R Core Team 2022) were used for graph generation.

## Results

### Comparing unit-specific antibiograms to the hospital-wide antibiogram

Figure 1 presents the weighted average differences in specific units (emergency department [ED], intensive care unit [ICU], transplant [TR] and units that are not ED, ICU, nor TR [nEIT]) when compared to a hospital-wide all-specimens antibiogram divided by gram-negative and gram-positive organisms.

**Figure 1.**
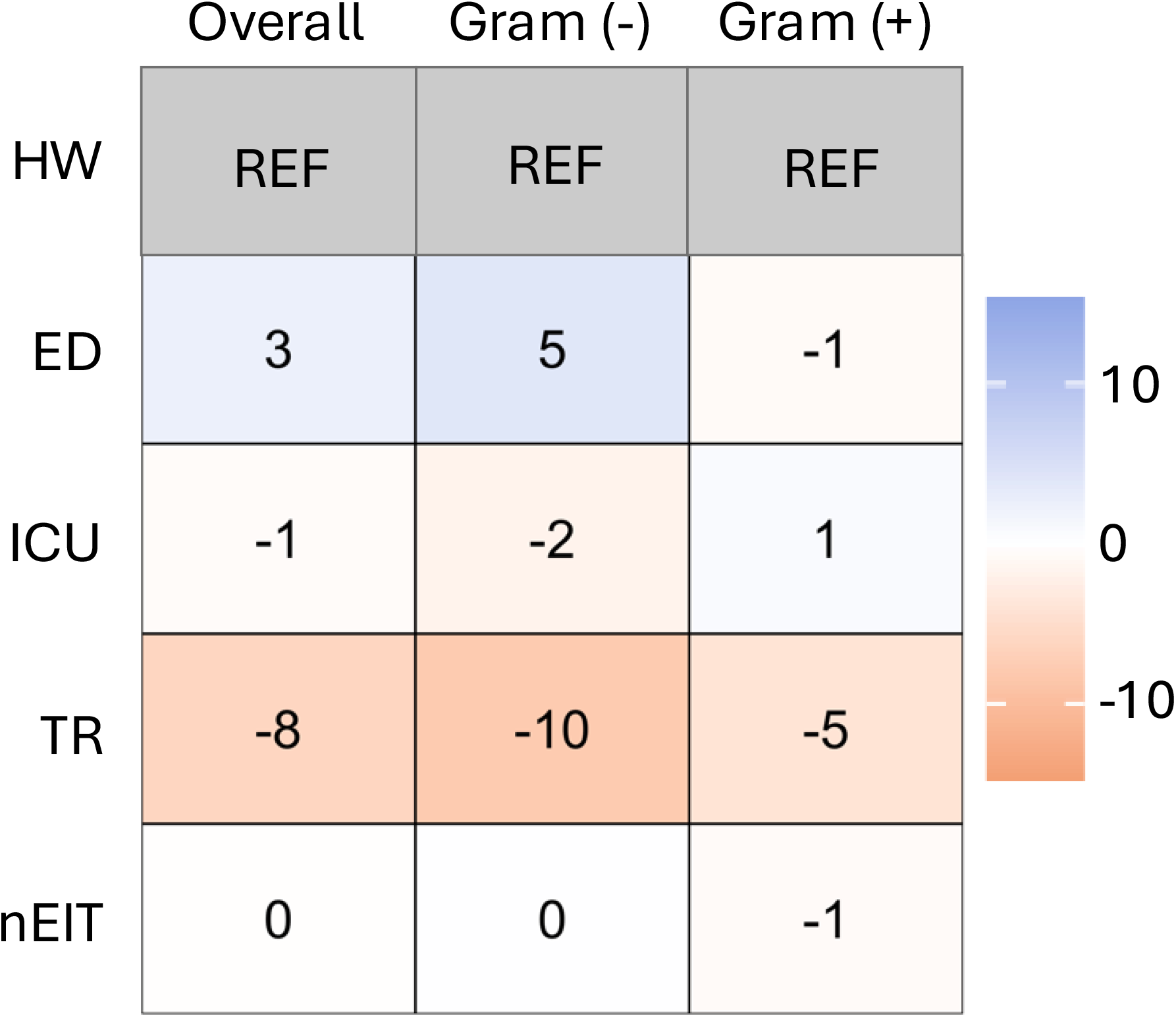
Comparison of unit-specific stratified antibiograms to the non-stratified hospital-wide antibiogram. Heat map displaying differences in susceptibility by weighted averages without CNST for specific units (emergency department [ED], intensive care unit [ICU], transplant [TR] and units that are not ED, ICU nor TR [nIET]) compared to hospital-wide (HW). Overall weighted average as well as Gram-negative and Gram-positive weighted averages are shown.

### Comparing specimen-stratified (syndromic) antibiograms within wards to ward-level antibiograms

Figure 2 provides heat maps showing the weighted average for specific specimens (blood, urine, resp and nBUR) among units: ED (Figure 2a), ICU (Figure 2b), TR (Figure 2c), and nEIT (Figure 2d).

**Figure 2.**
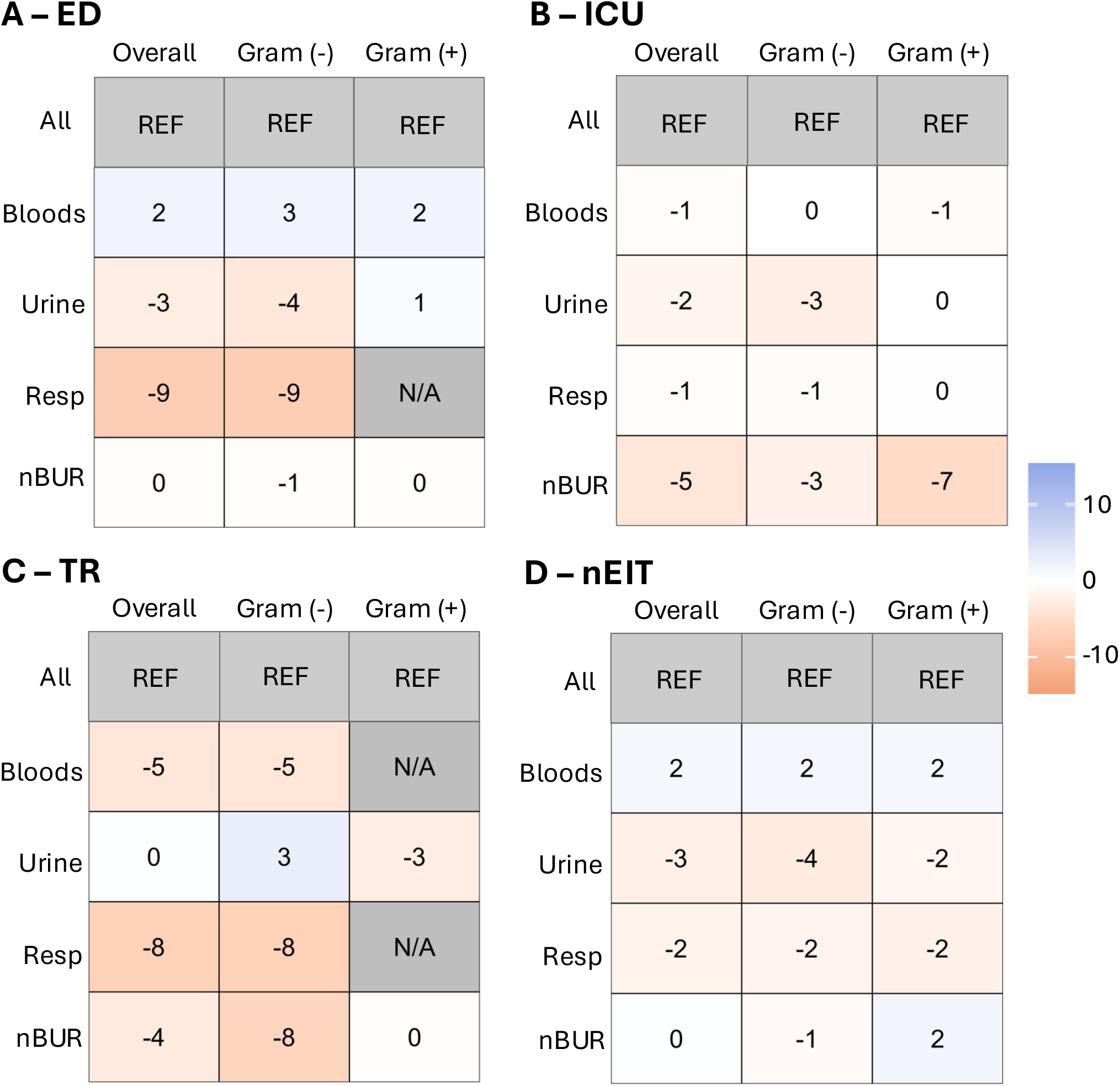
Comparison of specimen-stratified (syndromic) antibiograms within wards to the ward-level antibiogram. Heat map displaying the weighted average for specific specimen types (blood, urine, resp [resp] and non-blood/urine/resp [nBUR]) compared to all specimens (ALL) amongst different units: ED **(Figure 2a)**, ICU **(Figure 2b)**, TR **(Figure 2c)**, and units that are not ED, ICU nor TR (nIET; **Figure 2d)** are shown,. For Figure 2a-d, overall weighted average as well as Gram-negative and Gram-positive weighted average are shown.

### Summary of weighted average of the change in percent susceptibility between antibiograms

Figure 3 summaries the overall weighted averages of the change in percent susceptibility comparing different wards when using a wards-specific antibiograms as reference.

**Figure 3.**
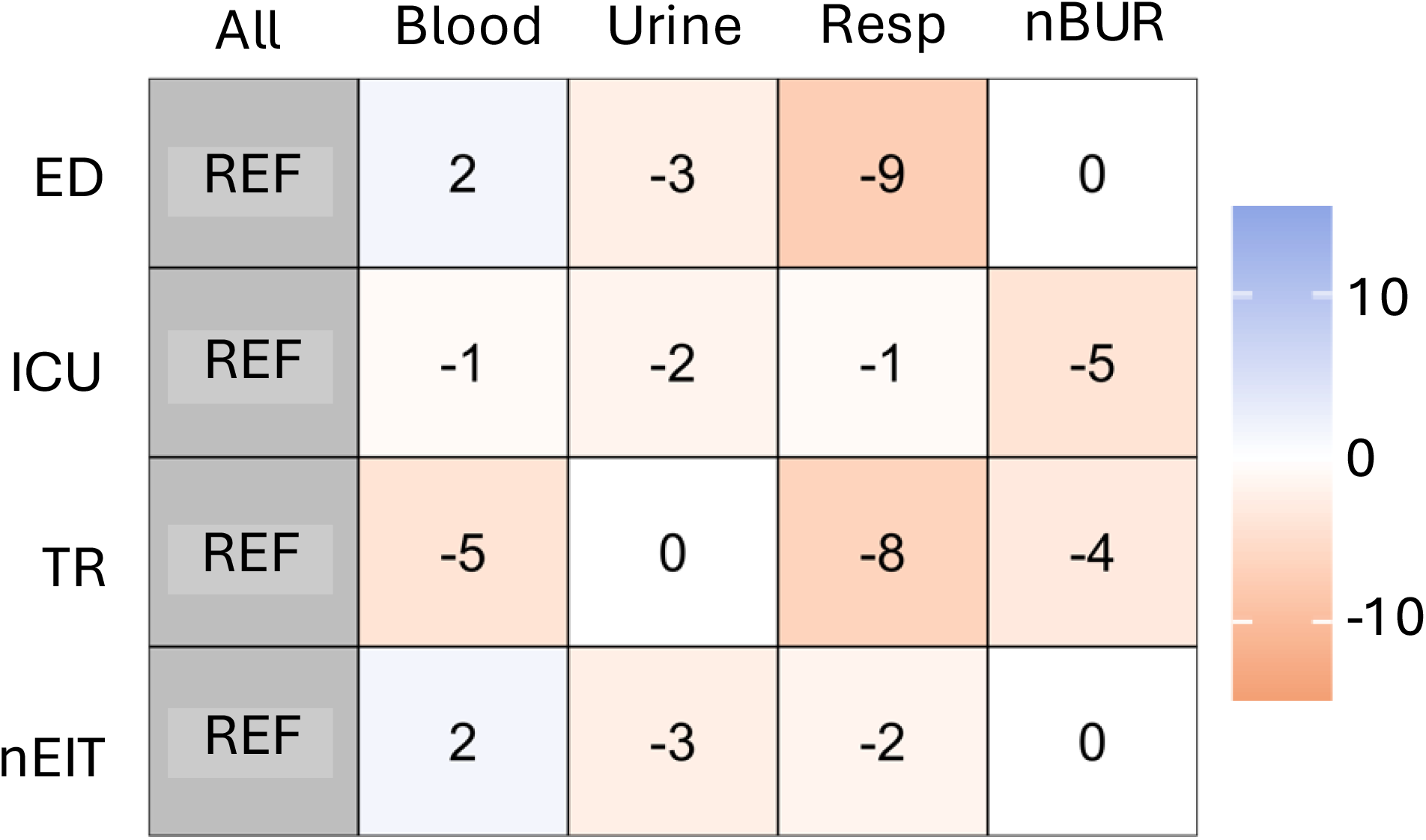
Summary of weighted averages of the change in percent susceptibility between antibiograms. Heat maps showing weighted average of the change in percent susceptibility between antibiograms for unit (emergency department [ED], intensive care unit [ICU], transplant [TR] and units that are not ED, ICU nor TR [nIET])/specimen (blood, urine, respiratory [resp], and specimens that are not blood, urine nor resp [nBUR]) combinations using unit-specific antibiograms as reference.

## Discussion

This study evaluated the weighted average of the change in percent susceptibility by organism in stratified antibiograms compared to non-stratified antibiograms. Differences reflect both differences in percent susceptibility of the organisms and differences in the abundance of the organism for that stratification’s demographic. In this study, only the organism abundance of the stratified antibiogram is taken into account, not the reference antibiogram.

Comparing unit-specific antibiograms through weighted average in Figures 1 showed that there is value-added to having unit-specific antibiograms particularly for TR units. Overall, the ED antibiogram showed increased susceptibility for all types of bacteria, especially Gram-negative bacteria. The opposite trend was revealed for TR where the TR antibiogram had reduced susceptibility compared to the non-stratified hospital-wide all-specimen antibiogram. When looking at the most prevalent organisms in Appendix 1, similar unit trends were seen in most organism-antimicrobial combinations. Some notable findings are that *E. coli* ciprofloxacin susceptibility in TR exhibited a large decrease in susceptibility at 31% and *E. faecium* vancomycin showed reduced susceptibility in TR of 13%.

Comparing the specimen-specific (syndromic) antibiograms stratfied by unit by weighted average to unit-specific antibiograms in Figure 2 highlights the value of this additional layer of stratification. Compared to the all-specimen ED stratified antibiogram (Figure 2a), respiratory isolates, though limited to *P. aeruginosa*, show a substantial decrease in susceptibility. Urine isolates also display an overall trend of reduced susceptibility, while blood isolates show an increase in susceptibility. Notable changes in susceptibility for blood isolates include increases for *E. coli* to ampicillin (23%), *P. mirabilis* to ampicillin (13%) and ciprofloxacin (17%), and for methicillin-susceptible *S. aureus* (9%). In urine isolates, there are significant reductions in susceptibility for *K. pneumoniae* to tobramycin (58%), *P. mirabilis* to ampicillin (42%) and tobramycin (27%), *K. oxytoca* to tobramycin (42%), *E. coli* to tobramycin (30%) and ampicillin (24%). Respiratory samples show a 10% reduction in susceptibility for *P. aeruginosa* to piperacillin/tazobactam and meropenem. Additionally, *E. coli* in nBUR specimens demonstrates reduced susceptibility to ampicillin (24%), piperacillin/tazobactam (20%), and amoxicillin/clavulanic acid (12%).

The same analysis was performed for ICU, as shown in Figure 2b, revealing slight reductions in susceptibility across all specimen types, with nBUR demonstrating the most significant relative reduction. Notable changes include reduced susceptibility of blood isolates of *E. coli* to ciprofloxacin (10%), ceftriaxone (8%), and ceftazidime (8%), while ampicillin (16%) and tobramycin (9%) showed increased susceptibility. In urine isolates of *E. coli*, susceptibility decreased for tobramycin (59%), ertapenem (30%), and ampicillin (22%), but increased for ceftriaxone (5%) and meropenem in *P. aeruginosa* (17%). Respiratory isolates exhibited reduced susceptibility of *E. coli* to tobramycin (26%), ampicillin (22%), amoxicillin/clavulanic acid (14%), piperacillin/tazobactam (12%), and ceftriaxone (8%) as well as reduced susceptibility of *K. pneumoniae* to amikacin (12%) and ceftriaxone (8%). For nBUR samples, susceptibility was reduced for meropenem in *P. aeruginosa* (19%) and vancomycin in *E. faecium* (11%), while susceptibility increased for tobramycin (8%) and ciprofloxacin (7%) in *E. coli*. For TR samples, as shown in Figure 2c, isolate numbers became more restricted. The urine subgroup antibiograms did not reveal any significant overall trends compared to the all-specimen TR antibiogram based on weighted average. However, there were observable trends of decreased susceptibility in blood, respiratory, and nBUR isolates compared to the hospital-wide respiratory antibiogram. On an individual organism basis (Appendix 4), blood *E. coli* exhibited reduced susceptibility to ciprofloxacin (12%), amikacin (11%), ceftriaxone (8%), and ceftazidime (8%). In urine isolates, *E. coli* and *K. pneumoniae* showed a significant reduction in susceptibility to tobramycin (31% and 23%, respectively), with a mild decrease observed in *E. faecium* for vancomycin (8%). Conversely, susceptibility increased in urine isolates for *P. aeruginosa* to gentamicin (21%), amikacin (19%), piperacillin/tazobactam (18%), ceftazidime (18%), and tobramycin (14%), as well as for *E. coli* to ceftriaxone (8%) Respiratory samples showed reduced susceptibility for *P. aeruginosa* to gentamicin (14%), ceftazidime (11%), and amikacin (10%). In nBUR samples, *P. aeruginosa* exhibited reduced susceptibility to amikacin (17%), gentamicin (15%), and tobramycin (15%).

Differences were also noted in units other than those described (nEIT, Figure 2d). Increased susceptibility in bloods were noted, while the urine and respiratory subgroup antibiograms showed decreased susceptibility compared to the all-specimens nEIT antibiogram. At the organism-drug level in Appendix 5, blood isolates show reduced susceptibility in *P. aeruginosa* for piperacillin/tazobactam (13%) and *K. pneumoniae* for ceftriaxone (9%), ceftazidime (9%), amoxicillin/clavulanic acid (7%) and piperacillin/tazobactam (7%), whereas increases in susceptibility were seen for *E. coli*-ampicillin (25%), *K. pneumoniae-*tobramycin (13%), *E. faecium-*vancomycin (12%), *E. coli-*tobramycin (9%), and increases in methicillin-susceptible *S. aureus* (5%). Urine isolates saw major reductions in susceptibility in *K. oxytoca-*tobramycin (88%), *K. pneumoniae-*tobramycin (49%), *P. mirabilis-*ampicillin (42%), *P. mirabilis-*amikacin (33%), and *E. coli-*tobramycin (29%). For respiratory isolates, notable susceptibility reductions were seen in *K. pneumoniae* for piperacillin/tazobactam (12%), ciprofloxacin (11%), tobramycin (11%), trimethoprim-sulfamethoxazole (10%) and amoxicillin/clavulanic acid (7%). A similar pattern was noted for *E. coli* with reductions in susceptibility seend in amoxicillin/clavulanic acid (17%), ampicillin (13%), piperacillin/tazobactam (9%), ceftriaxone (6%) and ceftazidime (6%) with increases seen for tobramycin (22%), ciprofloxacin (10%) and gentamicin (10%). Lastly, in nBUR samples, susceptibility reductions are seen for *E. coli* in ceftriaxone (11%), amoxicillin/clavulanic acid (10%) and piperacillin/tazobactam (9%), as well as *K. pneumoniae* in ciprofloxacin (9%), ertapenem (6%), gentamicin (5%) and amikacin (5%). Additionally, increased susceptibility was seen for *P. mirabilis* in amikacin (17%) and in methicillin-susceptible *S. lugdunensis* at 7%.

These results shown both visually through heat maps and numerically through weighted average differences in susceptibility suggest there is value-added in continuing to provide specimen-specific (syndromic) antibiograms stratified by unit to help best inform empiric therapy for specific populations. It is clear from our data that using a hospital-wide all-specimens antibiogram may both overcall and under call susceptibility leading to poor empiric antimicrobial choices. While others have recommended ICU-specific antibiograms [20,21] and ED-specific antibiograms [22], our data suggest value-added in further extending unit-specific data to include other units of interest such as TR units, and to further stratify unit-specific data by specimen type (syndrome) in order to identify nuanced changes in susceptibility that would otherwise go unnoticed in non-stratified antibiograms. The small numbers that may result from such sub-stratification should be recognized as a limitation with a caution that precision around those subgroup estimates of susceptibility may not be high. Collecting data from a larger time duration for those substrata may be helpful to obtain more precise estimates of susceptibility. It is recognized that this study was performed using data collected from a large urban area quaternary-care academic center and the results may not be generalizable to all. Laboratories and antimicrobial stewardship programs are encouraged to examine the specific needs of their own populations and to recognize the limitations of non-stratified antibiograms.

## Data Availability

Data not publicly available.

## Acknowledgements

Connie Xie completed data acquisition, completed all analyses, figures, and interpretations of results, and wrote the first draft of the manuscript. Samantha Martinez and Ceylon Simon completed data acquisition and reviewed and edited the manuscript. Susan Poutanen conceived and designed the study, oversaw the data acquisition, analyses and interpretation, and reviewed and revised the manuscript. All authors reviewed and approved the final version of the manuscript. The authors would like to thank Poolak Akhavan for her hard work in developing the original stratified antibiogram templates as well as Alyssa Loughborough, Melissa Kissoon, Bryn Hazlett, Shaista Anwer, Suresh Tharmaradinam, Isabella McNamara, and Pranav Tandon for their help in modifying and fine-tuning these templates.

## Abbreviations

ED: emergency department
ICU: intensive care unit TR transplant
nEIT: units that are not ED, ICU, nor TR
nBUR: specimens that are not blood, urine or respiratory

## Appendix Legends

**Appendix 1.**
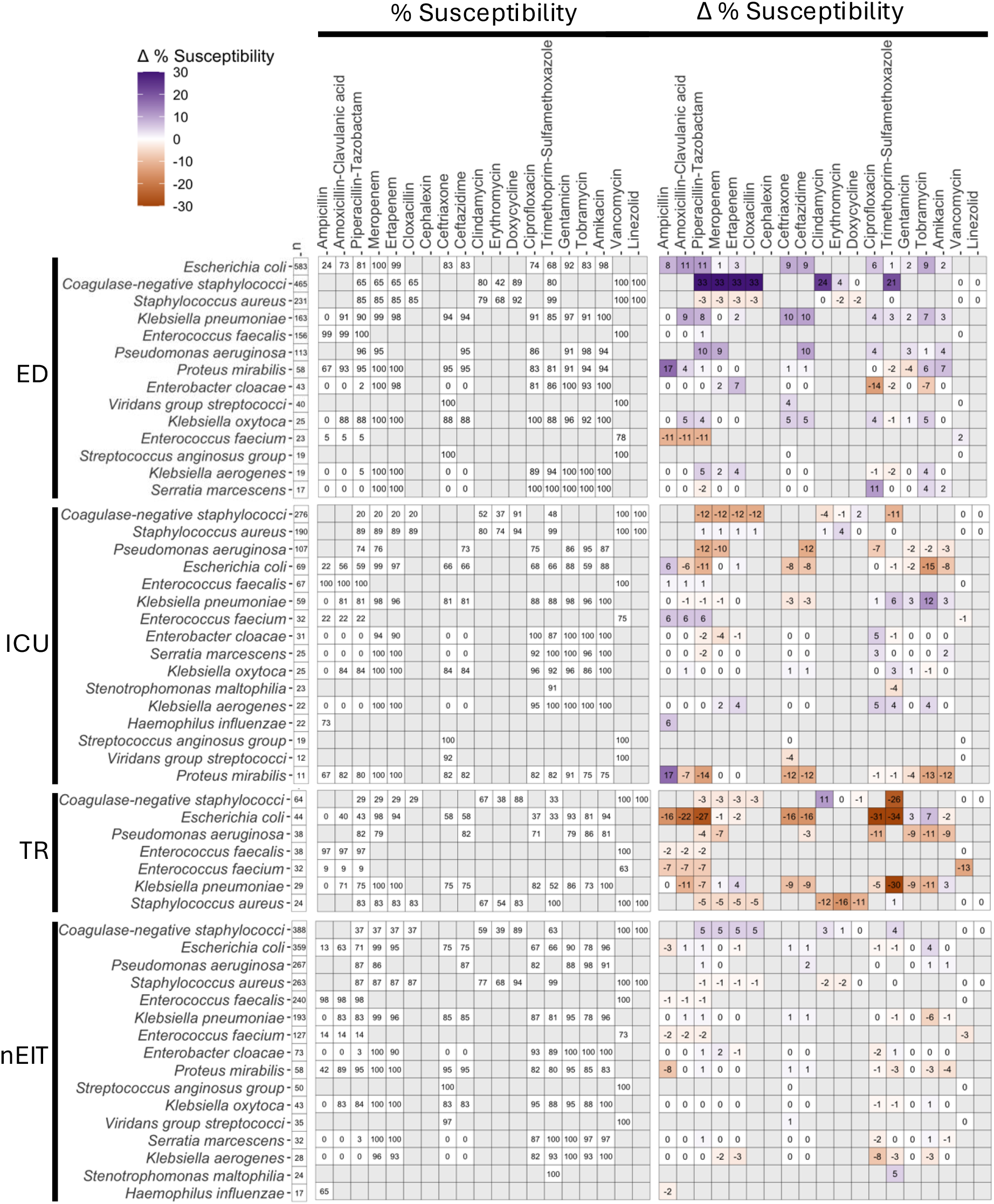
Heat map displaying differences in susceptibility percentages by individual organism/antimicrobial combinations for unit-specific (emergency department [ED], intensive care unit [ICU], transplant [TR] and units that are not ED, ICU nor TR [nIET]) all-specimens stratified antibiograms compared to the non-stratified hospital-wide all-specimens antibiogram.

**Appendix 2.**
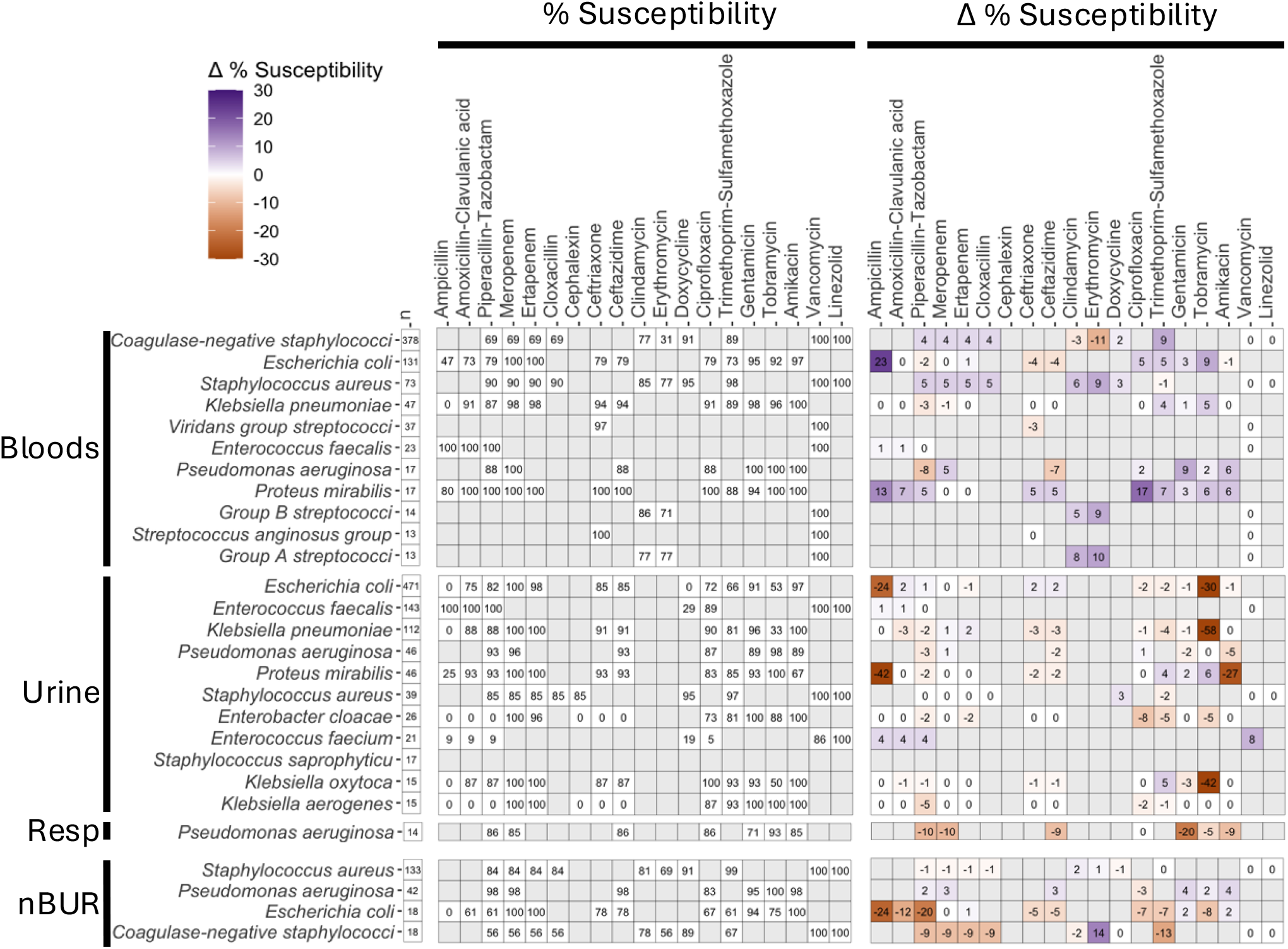
Heat map displaying differences in susceptibility percentages by individual organism/antimicrobial combinations for specimen-specific (blood, urine, respiratory [resp], and specimens that are not blood, urine nor resp [nBUR]) ED-only stratified antibiograms compared to the hospital-wide ED-only antibiogram.

**Appendix 3.**
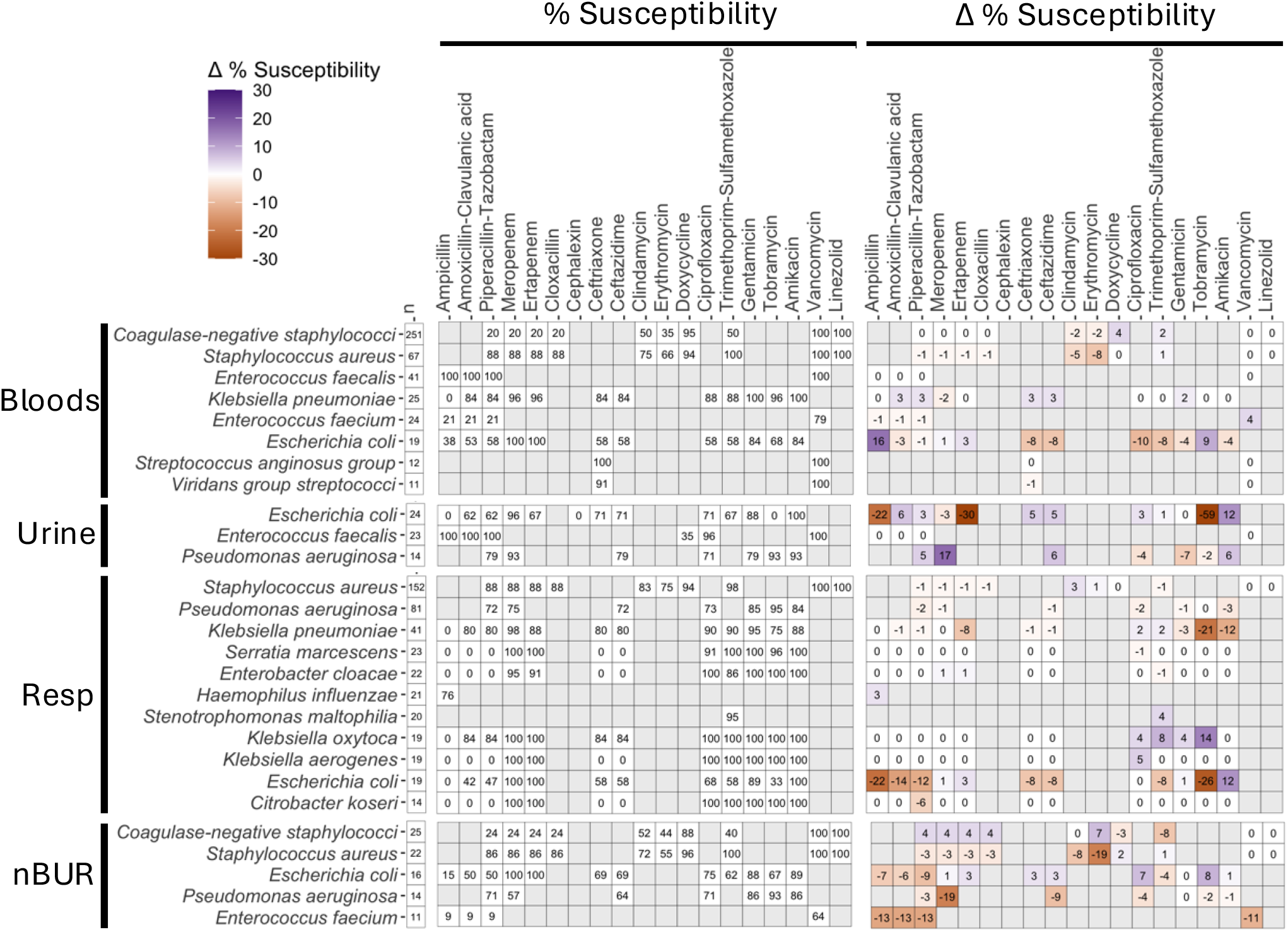
Heat map displaying differences in susceptibility percentages by individual organism/antimicrobial combinations for specimen-specific (blood, urine, respiratory [resp], and specimens that are not blood, urine nor resp [nBUR]) ICU-only stratified antibiograms compared to the hospital-wide ICU-only antibiogram.

**Appendix 4.**
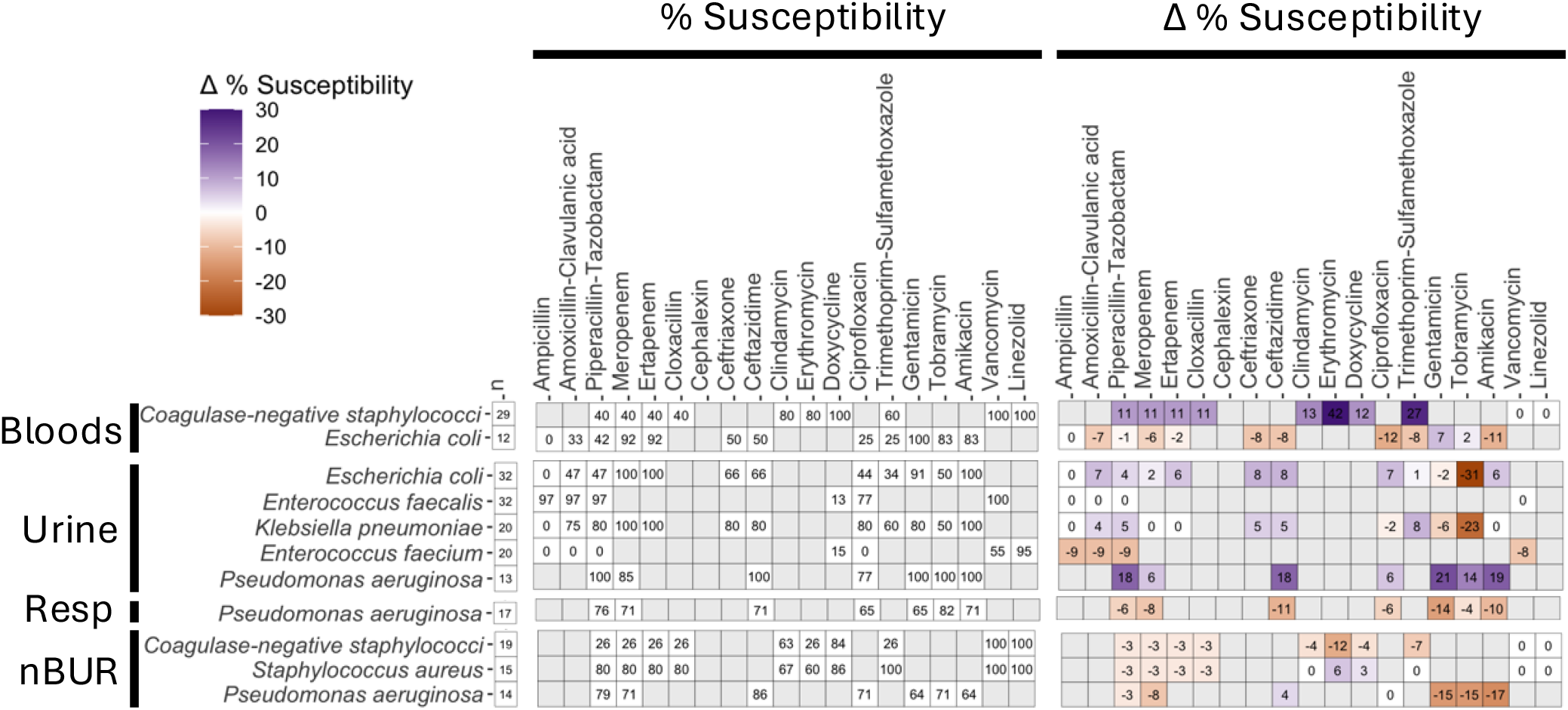
Heat map displaying differences in susceptibility percentages by individual organism/antimicrobial combinations for specimen-specific (blood, urine, respiratory [resp], and specimens that are not blood, urine nor resp [nBUR]) TR-only stratified antibiograms compared to the hospital-wide TR-only antibiogram.

**Appendix 5.**
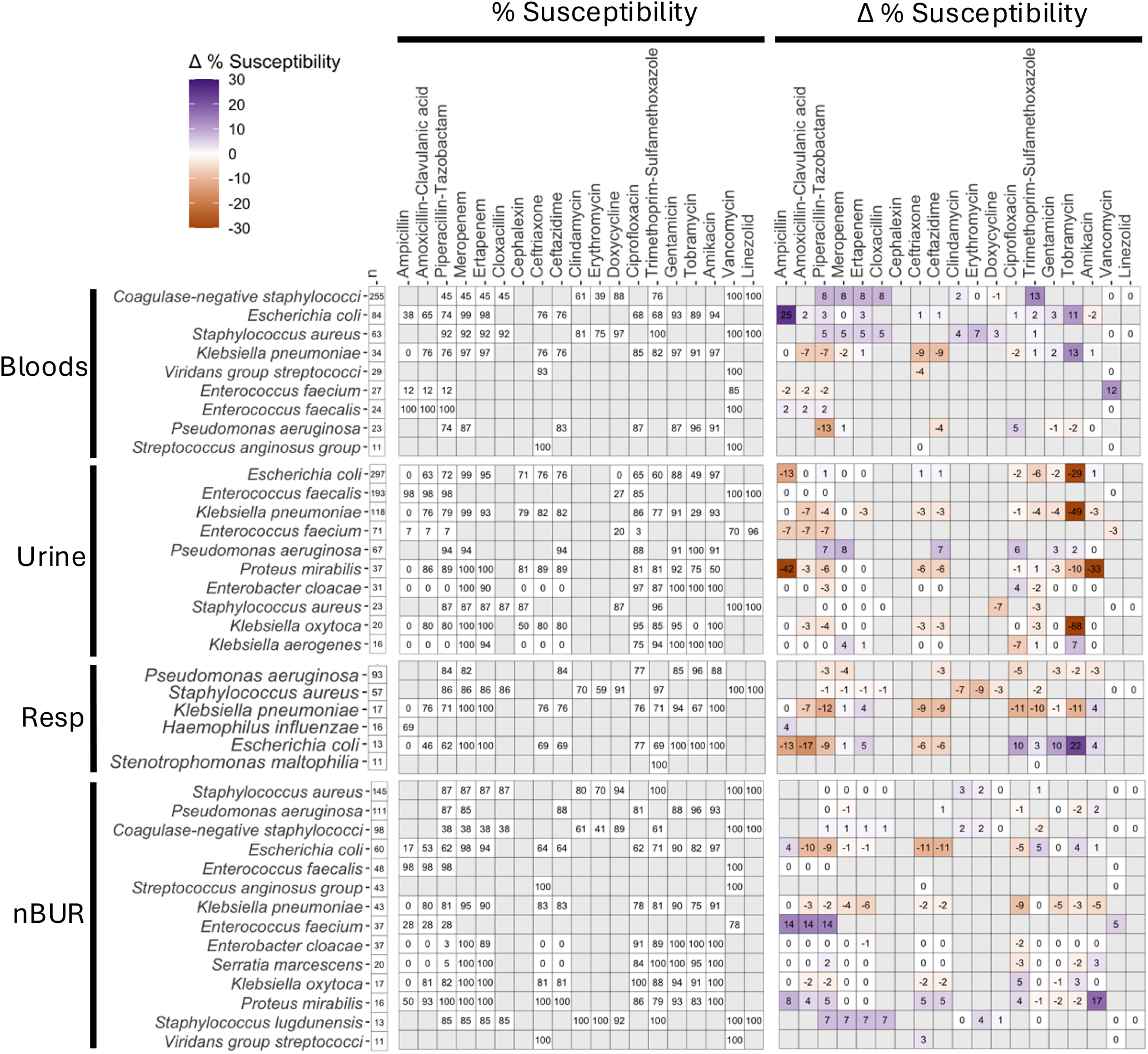
Heat map displaying differences in susceptibility percentages by individual organism/antimicrobial combinations for specimen-specific (blood, urine, respiratory [resp], and specimens that are not blood, urine nor resp [nBUR]) nEIT-only stratified antibiograms compared to the hospital-wide nEIT-only antibiogram.

